# Stochastic Lanczos estimation of genomic variance components for linear mixed-effects models

**DOI:** 10.1101/607168

**Authors:** Richard Border, Stephen Becker

## Abstract

**Background:** Linear mixed-effects models (LMM) are a leading method in conducting genome-wide association studies (GWAS) but require residual maximum likelihood (REML) estimation of variance components, which is computationally demanding. Previous work has reduced the computational burden of variance component estimation by replacing direct matrix operations with iterative and stochastic methods and by employing loose tolerances to limit the number of iterations in the REML optimization procedure. Here, we introduce two novel algorithms, *stochastic Lanczos derivative-free REML* (`SLDF_REML`) and *Lanczos first-order Monte Carlo REML* (`L_FOMC_REML`), that exploit problem structure via the principle of Krylov subspace shift-invariance to speed computation beyond existing methods. Both novel algorithms only require a single round of computation involving iterative matrix operations, after which their respective objectives can be repeatedly evaluated using vector operations. Further, in contrast to existing stochastic methods, `SLDF_REML` can exploit precomputed genomic relatedness matrices (GRMs), when available, to further speed computation.

**Results:** Results of numerical experiments are congruent with theory and demonstrate that interpreted-language implementations of both algorithms match or exceed existing compiled-language software packages in speed, accuracy, and flexibility.

**Conclusions:** Both the `SLDF_REML` and `L_FOMC_REML` algorithms outperform existing methods for REML estimation of variance components for LMM and are suitable for incorporation into existing GWAS LMM software implementations.

Full list of author information is available at the end of the article

## Background

Linear mixed-effects modeling (LMM) is a leading methodology employed in genome-wide association studies (GWAS) of complex traits in humans, offering the dual benefits of controlling for population stratification while permitting the inclusion of data from related individuals [1]. However, the implementation of LMM comes at the cost of increased computational burden relative to ordinary least-squares regression, particularly in performing residual maximum likelihood (REML) estimation of genomic variance components. Conventional REML algorithms require multiple 𝒪 (*n*^3^) or 𝒪 (*mn*^2^) matrix operations, where *m* and *n* are the numbers of markers and individuals, respectively, rendering them infeasible for large biobank scale data sets. As a result, the problem of increasing the computational efficiency of REML estimation of genomic variance components has generated considerable research activity [2–6].

In the case of the standard two variance component model (1), the estimation of which is the focus of the current research, previous efforts toward increasing computational efficiency fit into two primary categories: 1., reducing the number of cubic time complexity matrix operations needed to achieve convergence; and 2., substituting stochastic and iterative matrix operations for deterministic, direct methods to obtain procedures with quadratic time complexity. The first approach is embodied by the methods implemented in the FaST-LMM and GEMMA packages [2, 4], which take advantage of the fact that the genetic relatedness matrix (GRM) and identity matrix comprising the covariance structure are simultaneously diagonalizable. As a result, after performing a single spectral decomposition of the GRM and a small number of matrix-vector multiplications, the REML criterion and its gradient and Hessian can be repeatedly evaluated using only vector operations. The second approach is exemplified by the popular BOLT-LMM software [5, 6], which avoids all cubic operations by solving linear systems via the method of conjugate gradients (CG) and employing stochastic trace estimators in place of deterministic computations.

In the current research, we propose two algorithms, stochastic Lanczos derivative-free residual maximum likelihood (`SLDF_REML`; Algorithm 3) and Lanczos first-order Monte Carlo residual maximum likelihood (`L_FOMC_REML`; Algorithm 4), that combine features of both approaches (Figure 1). Here, we translate the simultaneous diagonalizability of the heritable and nonheritable components of the covariance structure to stochastic and iterative methods via the principle of Krylov subspace shift-invariance. As a result, we only need to compute the costliest portions of the objective function once (via stochastic/iterative methods), computing all subsequent iterations of the REML optimization problem only using vector operations. We develop the theory underlying these methods and demonstrate their performance relative to previous methods via numerical experiment.

**Figure 1:**
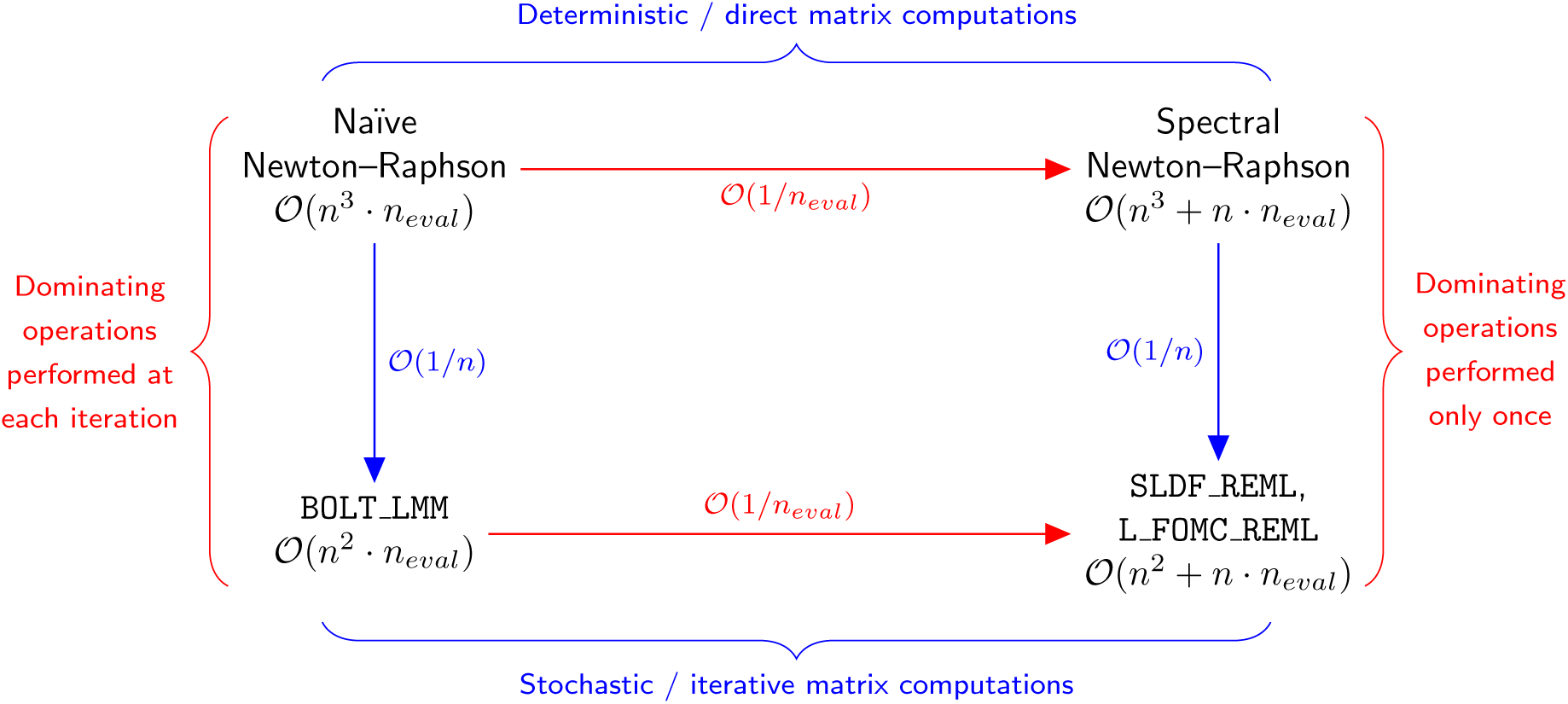
Time complexity analogies with respect to existing and proposed methods. Heuristically, the novel algorithms (bottom right) are to the stochastic, iterative algorithm implemented in the BOLT-LMM software [5, 6] (bottom left) as the direct methods exploiting the shifted structure of the the two component genomic variance component model (1) (e.g., FaST-LMM and GEMMA [2, 4]; top right) are to standard direct methods (top left). For simplicity, we assume here that the number of markers is equal to the number of observations and omit low-order terms related to the spectral conditioning of the covariance structure and the number of random vectors generated by the stochastic methods; further details are provided in Table 1. *n*_*eval*_ denotes the number of objective function evaluations needed to achieve convergence.

**Figure 2:**
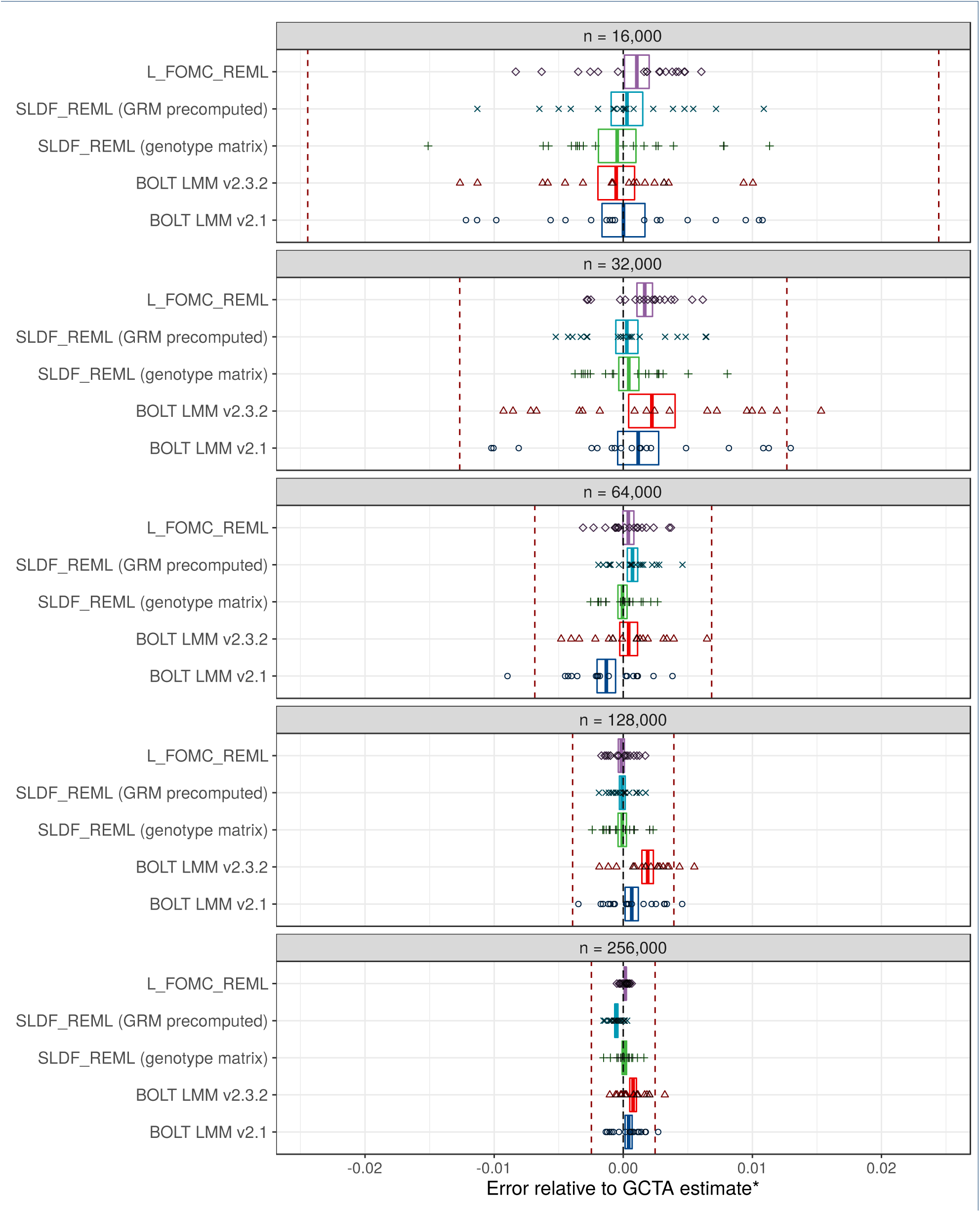
Accuracy results. Comparison of heritability estimates for height generated by BOLT-LMM versions 2.1 and 2.3.2, `SLDF_REML`, and `L_FOMC_REML` versus those generated by the deterministic algorithm implemented in the GCTA software package^∗^[3], for varying sub-samples of 16,000 to 256,000 unrelated European-ancestry UK Biobank participants. Data are comprised of twenty independent replications per condition. Red dashed lines indicate standard errors of GCTA estimate. Points represent individual observations whereas boxes indicate the 95% confidence intervals for the trimmed mean estimate after a Bonferroni correction for 25 comparisons. The bias evidenced by the BOLT-LMM estimators is likely due to the combination of performing a small number of secant iterations with fixed start values and loose tolerances for determining convergence. ^∗^For *n*=256,000, memory requirements prohibited the use of GCTA, so we instead averaged ten estimates generated by the high-accuracy stochastic estimator implemented in BOLT-REML [29] (standard errors were 6.32e-5 and 2.45e-7 for the mean REML heritability estimate and its standard error, respectively).

**Figure 3:**
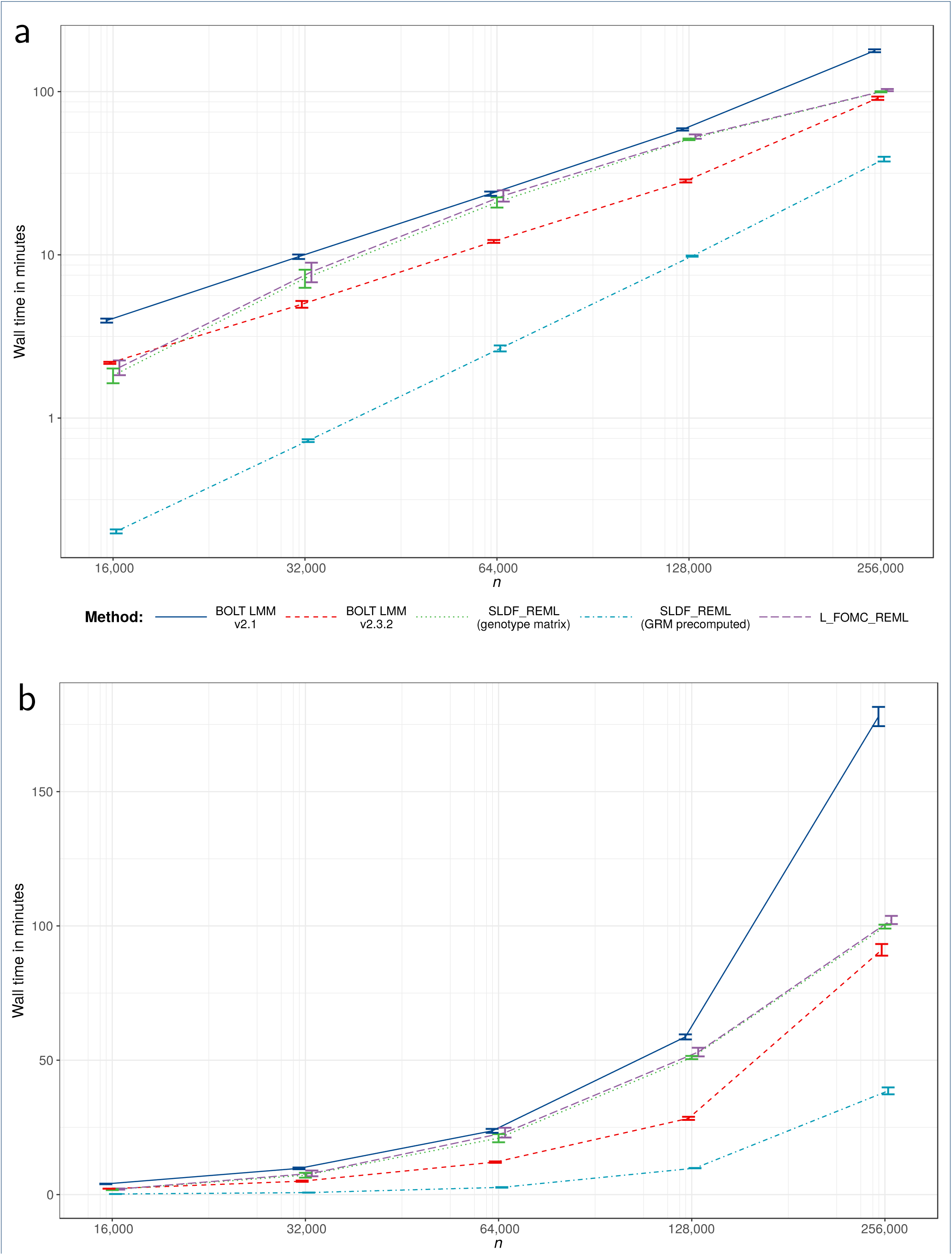
Timing results. Trimmed mean wall clock time across twenty replications for per condition on the log10 scale (a) and natural scale (b). Error bars reflect per condition standard errors and lines connect per condition trimmed means.

## Method

We consider the two component genomic variance components model commonly employed in LMM association studies [1], which is of the form

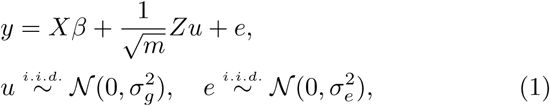

where *y* is a measured phenotype, the *c* ≪ *n* columns of *X* ∈ ℝ^*n×c*^ are covariates (including an intercept term) with corresponding fixed effects *β*, and *Z* ∈ ℝ^*n×m*^ is a matrix of *n* individuals’ standardized genotypes at *m* loci. Without loss of generality, we assume that *X* has full column rank; in the case of numerical rank deficiency we can simply replace *X* by the optimal full rank approximation generated by its economy singular value decomposition or rank revealing QR decomposition. The latent genetics effects *u* ∈ ℝ^*m*^ and residuals *e* ∈ ℝ ^*n*^ are random variables with distributions parametrized by the heritable and non-heritable variance components, 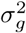 and 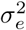, respectively. The REML criterion corresponds to the marginal likelihood of 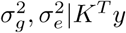, where *K*^*T*^ projects to an (*n − c*)dimensional subspace orthogonal to the covariate vectors such that the null space of *K*^*T*^ is exactly the column space of *X* [7]. In other words *K*^*T*^ : ℝ^*n*^ *→* 𝒮⊂ ℝ^*n-c*^ such that ℝ^*n*^ = 𝒮 ⊕ col *X*. The transformed random variable *K*^*T*^ *y* has the marginal distribution 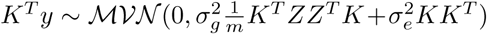, which we reparametrize as 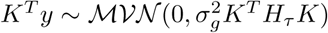, where

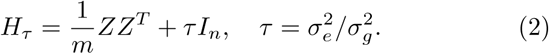

Here,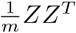, which indicates the average covariance between individuals’ standardized genotypes, is often referred to as the *genomic relatedness matrix* (GRM). The *REML criterion*, or marginal log likelihood, can be expressed as a function of *τ* :

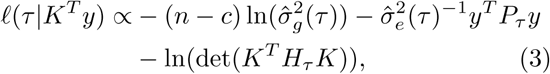

where *P*_*τ*_ = *K*(*K*^*T*^ *H*_*τ*_ *K*)^*-*1^*K*^*T*^, and, as implied by the REML first-order (stationarity) conditions, 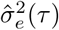 is the expected residual variance component given *τ* and 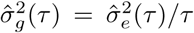 [7, 8]. In practice, *K* is never explicitly formed.

**Table 1:**
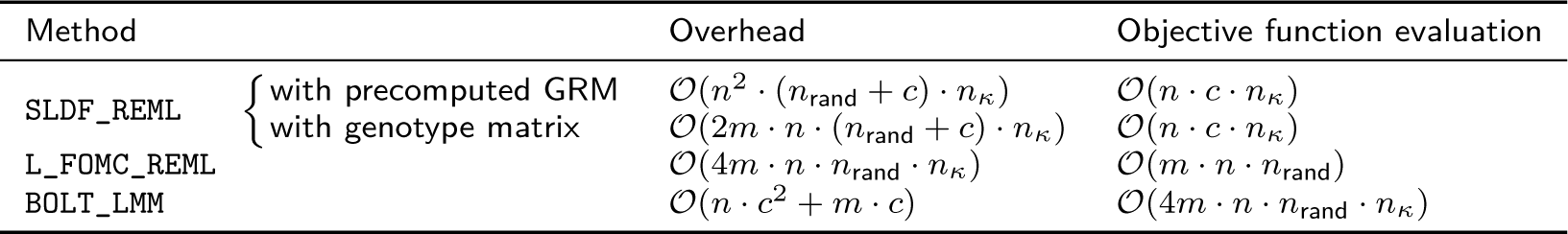
Time complexity of stochastic algorithms. *n* denotes the number of individuals, *m* the number of markers, and *c* the number of covariates. *n*_rand_ indicates the number of random probing vectors and is fixed at 15 in all numerical experiments. *n*_*κ*_ reflects the number of conjugate gradient iterations required to achieve convergence at a specified tolerance and can be bounded in terms of the spectral condition number of *H*_0_. As noted in [6], implicit preconditioning of *H*_0_ can be achieved by including the first few right singular vectors of the genotype matrix (or eigenvectors of the GRM) as covariates.

Naïve procedures for maximizing the REML criterion require evaluating (3) or its derivatives at each iteration of the optimization procedure. Previous methods either reduce the number of necessary cubic time complexity operations to one by exploiting problem structure, or subsitute quadratic time complexity iterative and stochastic matrix operations for direct computations (Figure 1). Here, we unify these approaches via the principle of Krylov subspace shift invariance to achieve methods that only require a single iteration of quadratic time complexity operations.

In what follows, we first present a brief survey of the Lanczos process, its applications to families of shifted linear systems, and its use in constructing Gaussian quadratures for spectral matrix functions. We assume familiarity with the method of conjugate gradients and Gaussian quadrature; if not, see [9] and [10], respectively. We present methods toward the goal of efficiently evaluating the quadratic form and log-determinant terms appearing in the REML criterion (3). We then present the details of the `SLDF_REML` and `L_FOMC_REML` algorithms, both of which exploit problem structure via Lanczos process-based methods in order to speed computation. Finally, we derive expressions for the computational complexity of the present algorithms, which we confirm via numerical experiment.

### Preliminaries

The notation in this section is self-contained. Our presentation borrows from the literature extensively; further details on the (block) Lanczos procedure [9, 11], conjugate gradients for shifted linear systems [12, 13], stochastic trace estimation [14, 15], and stochastic Lanczos quadrature [16–18] are suggested in the bibliography.

### Krylov subspaces

Consider a symmetric positive-definite matrix *A* and nonzero vector *b*. Define the *m*^*th*^ Krylov subspace by the span of the first *m -* 1 monomials in *A* applied to *b*; that is, 𝒦_*m*_(*A, b*) = span {*A*^*k*^*b* : *k* = 0, *…*, *m -* 1}. Krylov subspaces are *shift invariant*—i.e., for real numbers *σ*, we have 𝒦_*m*_(*A, b*) = 𝒦_*m*_(*A* + *σI, b*).

### The Lanczos procedure

The Lanczos procedure generates the decomposition *AU*_*m*_ = *U*_*m*_*T*_*m*_, where the columns *u*_1_, …, *u*_*m*_ of *U*_*m*_ form an orthonormal basis for 𝒦_*m*_(*A, b*) and the *Jacobi matrices T*_*m*_ ∈ ℝ^*m×m*^ are symmetric tridiagonal. Choosing *u*_1_ = *b*/ ‖*b*‖, successive columns are uniquely determined by the sequence of Lanczos polynomials 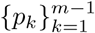 such that each *u*_*k*_ = *p*_*k-*1_(*A*)*u*_1_ and each *p*_*k*_ is the characteristic polynomial of Jacobi matrix *T*_*k*_ consisting of the first *k* rows and columns of *T*_*m*_. The Lanczos procedure is equivalent to the well-known method of conjugate gradients (CG) for solving the linear system *Ax* = *b* in that the *m*^*th*^ step CG approximate solution *x*^(*m*)^ is obtained from the above decomposition using only vector operations (see Algorithm 1).

The number of steps *m* prior to termination corresponds to the number of CG iterations need to bound the norm of the residual below a specified tolerance: ‖*Ax*^(*m*)^–*b* ‖ < ϵ. The rate of convergence depends on the spectral properties of *A* and can be controlled in terms of the spectral condition number *κ*(*A*). In the present application, the fact that all complex traits of interest generally have a non-trivial non-heritable component results in well-conditioned systems [5, 19].

### Solving families of shifted linear systems

Having applied the Lanczos process to the *seed system Ax* = *b*, shift-invariance can be exploited to obtain the *m*^*th*^ step CG approximate solution 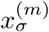 to the *shifted linear system A*_*σ*_*x*_*σ*_ = (*A* + *σI*)*x*_*σ*_ = *b*, only using vector operations [12]. It can be shown that any positive shift by *σ ≥* 0 improves the rate of convergence such that 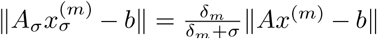, where *δ*_*m*_ *>* 0 is the *m*^*th*^ diagonal element of the Lanczos Jacobi matrix corresponding to 𝒦_*m*_(*A, b*).

### Lanczos polynomials and Gaussian quadrature

Additionally, the Lanczos polynomials comprise a sequence of orthogonal polynomials with respect to the *spectral measure*

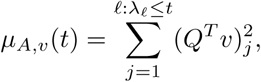

#### Algorithm 1

Lanczos conjugate gradients solver for shifted systems (`L_Solve`)

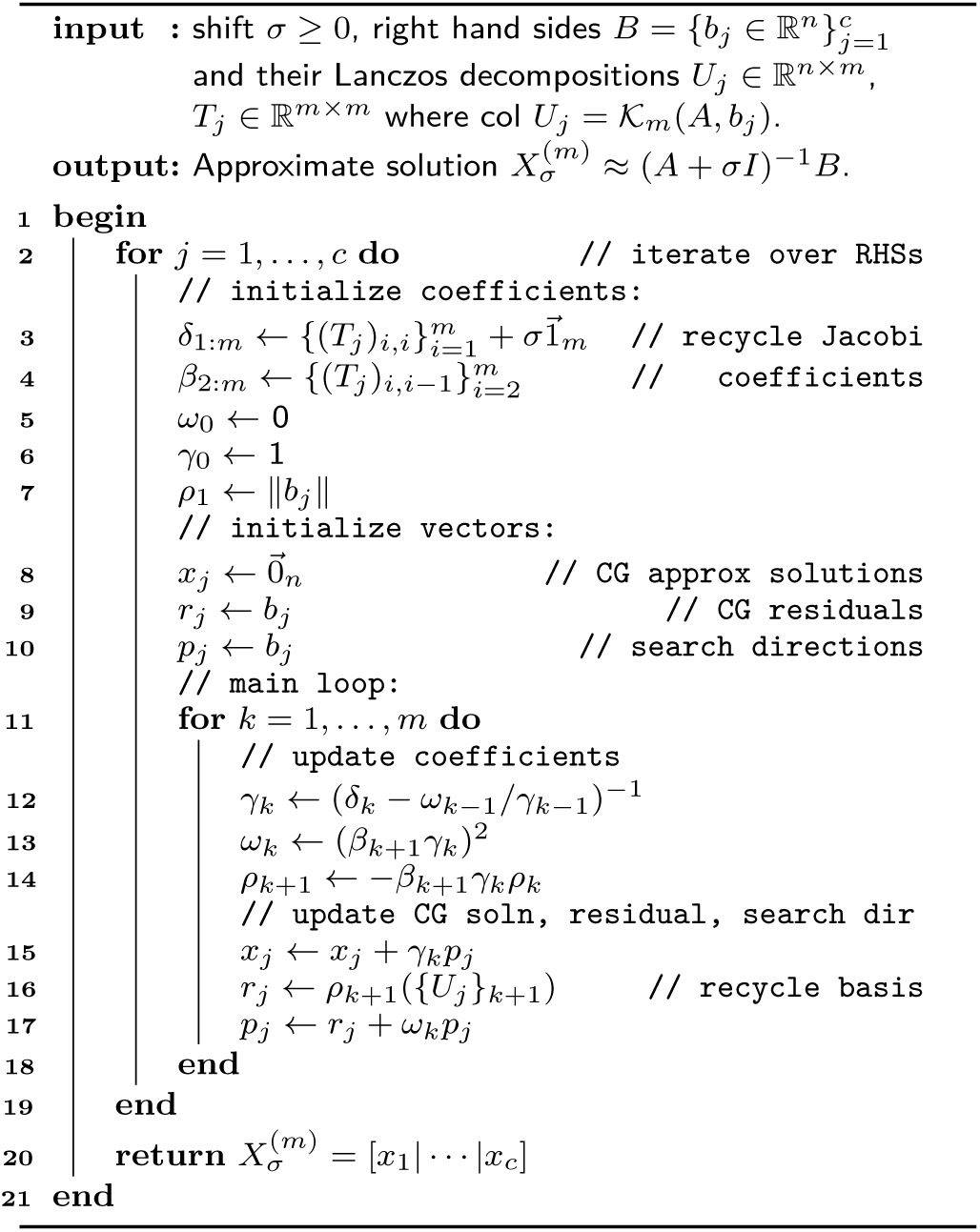

where *A* = *Q*Λ*Q*^*T*^ is the spectral decomposition [16, 17]. Quadratic forms *v*^*T*^ *f* (*A*)*v* involving *spectral functions f* (*A*) = *Qf* (Λ)*Q*^*T*^, e.g., for the matrix loga-rithm, 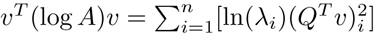, can be written as Riemann–Stieltjes integrals of the form

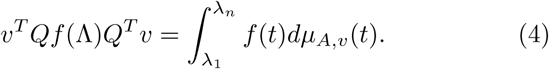

The Lanczos decomposition *AU*_*m*_ = *U*_*m*_*T*_*m*_ generates the weights and nodes for an *m*-point Gaussian quadrature approximating the above integral. Denoting the spectral decomposition of the *j*^*th*^ Jacobi matrix 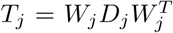 for *j* = 1, …, *m*, we approximate (4) as

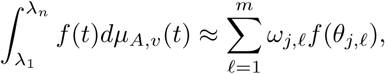

where *θ*_*j,ℓ*_ = *{D*_*j*_*}*_*ℓ,ℓ*_ and 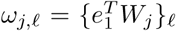. As *m* here corresponds to the number of CG iterations needed to ensure that ‖*Ax*^(*m*)^ –*v*‖ is smaller than a specified tolerance, the tridiagonal Jacobi matrices are small and calculating their spectral decompositions is computationally trivial

#### Algorithm 2

Stochastic Lanczos quadrature approximate log determinant of shifted systems (`SLQ_LDet`)

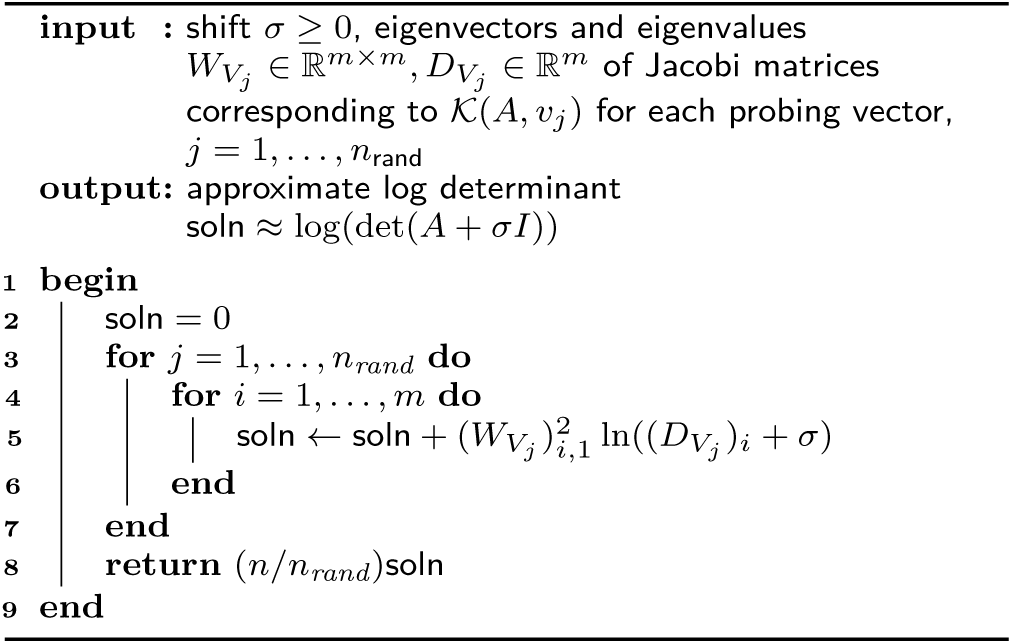

### Stochastic Lanczos quadrature

*Stochastic Lanczos quadrature* (SLQ) combines the above quadrature formulation with Hutchinson-type stochastic trace estimators [16]. Such estimators approximate the trace of a matri *H ∈ℝ*^*n*×*n*^ by a weighted sum of quadratic forms tr 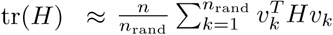 for normalized, suitably distributed i.i.d. random *probing vectors 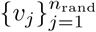* [14]. The SLQ approximate trace of a spectral function of a matrix, tr(*f* (*A*)), is then

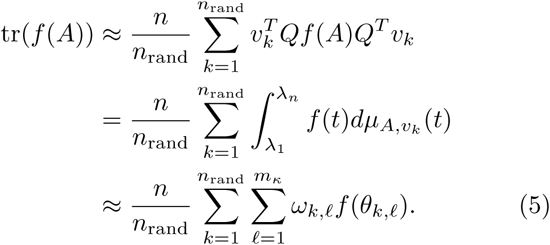

Whereas the number of probing vectors *n*_rand_ is chosen *a priori*, the number quadrature nodes *m*_κ_ corresponds to the number of conjugate gradient iterations needed to ensure *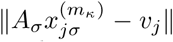* is less than a specified tolerance for each *j* = 1, …, *n*_rand_.

### SLQ and shift invariance

For a fixed probing vector *v*_*i*_, we can exploit the shift invariance of 𝒦_*m*_(*A, v*_*i*_) to efficiently update Gaussian quadrature generated by the corresponding Lanczos decomposition *AU*_*m*_ = *U*_*m*_*T*_*m*_. Again denoting the spectral decomposition of the Jacobi matrix 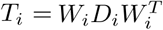, the Lanczos decomposition of the shifted system is simply 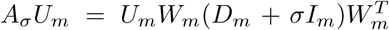 Thus, given the approximation (5) for tr(*f*(*A*)), we can efficiently compute an approximation of tr (*f(A*_*σ*_)) for any σ> 0.In Algorithm 2 we implement a method of tr (*f(A*_*σ*_)) in 𝒪 (*n*_rand_) operations given the spectral decompositions of the Jacobi matrices corresponding to 𝒦_*m*_*(A, v*_*j*_) for probing vectors 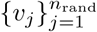.

### Block methods

For multiple right hand sides *B* = [*b*_1_| … |*b*_*c*_], the Lanczos procedure can be generalized to the *block Krylov subspace 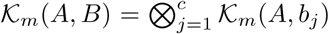*, result-ing in a collection of Lanczos decompositions *AU*_*j*_ =*U*_*j*_*T*_*j*_ such that *{U*_*j*_ *}*_1_ = *b /*‖*b* ‖ for *j* = 1, …, *c*. This process is equivalent to block CG methods in that the Jacobi matrices can again be used to generate an approximate solution *X*^(*m*)^ to the matrix equation *AX*^(*m*)^ = *B*. We provide an implementation of the block Lanczos procedure in `L_Seed` [20], employing a conservative convergence criterion defined in terms of the magnitude of the (1, 2) operator norm of the residual 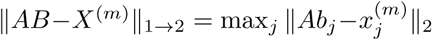 Compared to performing *c* separate Lanczos procedures with respect to 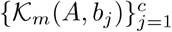, block Lanczos with respect to 𝒦_*m*_ *(A,B)* with *B* = [*b*_1_| … |*b*_*c*_], produces the same result (for a fixed number of steps). However, block Lanczos employs BLAS-3 operations and is thus more performant, especially when implemented on top of parallelized linear algebra subroutines.

### A derivative-free REML algorithm

We propose the stochastic Lanczos derivative-free residual maximum likelihood algorithm (`SLDF_REML;` Algorithm 3), a method for efficiently and repeatedly evaluating the REML criterion, which is then subject to a zeroth-order optimization scheme. To achieve this goal, we first identify the parameter space of interest with a family of shifted linear systems. We then develop a scheme for evaluating the quadratic form *y*^*T*^*P*_*τ*_*y* and log determinant ln (det (*K*^*T*^*H*_*τ*_*K))* terms in the REML criterion (3) that use the previously discussed Lanczos methods to exploit this shifted structure. Specifically, after obtaining a collection of Lanczos decompositions, we can repeatedly solve the linear systems involved in the quadratic form term via Lanczos conjugate gradients and approximate the log determinant term via stochastic Lanczos quadrature.

### The parameter space as shifted linear systems

Given a range of possible values of the *standardized genetic variance component*, or *heritability*,

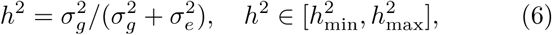

we set 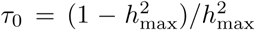 and define *H* _0_ = *H*_*τ0*_, noting that for all 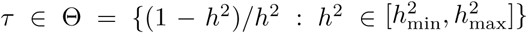 the spectral condition number of *H*_*τ*_ will be less than that of *H*_0_ as the identity component of *H*_*τ*_ will only increase. Further, we have now identified elements of our parameter space τ ∈Θ with the family of shifted linear systems

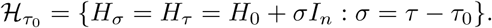

For any vector *v* for which we have computed the Lanczos decomposition *H*_0_*U* = *UT* with the first column of *U* equal to *v*/‖*v*‖, we can use Algorithm 1 to obtain the CG approximate solution 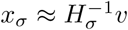 for all σ ≥0 in *O (n)* operations.

### The quadratic form

Directly evaluating the quadratic form

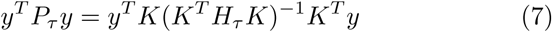

is computationally demanding and is typically avoided in direct estimation methods [7, 8]. Writing the complete QR decomposition of the covariate matrix *X* = [*Q*_*X*_|*Q*_*X ⊥*_] *R* allows us to define 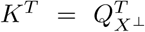 noting that substituting 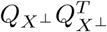 for *K*^*T*^ preserves the value of (7). 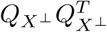 is equivalent to the orthogorthogonal projection operator 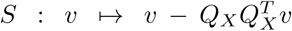 which admits an efficient implicit construction and is computed in *O*(*nc*^2^) operations via the *economy* QR decomposition *X* = *Q*_*X*_*R*_*X*_. Then, reexpressing (7) as *y*^*T*^ *S*(*SH*_*τ*_*S*)^†^*Sy*, we can use the Lanczos process to construct an orthonormal basis and corresponding Jacobi matrix for the Krylov subspace 𝒦(*SH*_0_*S, Sy*). We can then obtain the CG approximation of *y*^*T*^ *S*(*SH*_*σ*_*S*)^-1^*Sy* using vector operations as, for any shift σ, we have *y*^*T*^ *S*(*SH*_*σ*_*S*)^†^*Sy* = *y*^*T*^ *S*(*SH*_0_*S*+_σ_*In*)^-1^*Sy* (see Lemma 1 in Appendix for proof)

### The log determinant

We use an equivalent formulation [7, 21] of the term ln(det(*K*^*T*^*H*_*τ*_*K*)), rewriting it as

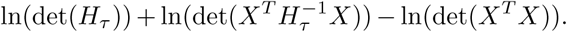

The det(*X*^*T*^*X*) term is constant with respect to τ and can be disregarded. For *c ≪ n*, det 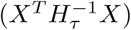 is computationally trivial via direct methods given 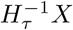, which we can compute for all parameter values of interest in *O*(*n*) operations having first applied the block Lanczos process with respect to 𝒦(*H*_0_,*X*). Computing the block Lanczos decomposition corresponding to 𝒦(*H*_0_,*X*), which is only performed once, unfortunately scales with the number of covariates *c*, a disadvantage not shared by our second algorithm (Algorithm 4). The remaining term, ln(det(*H*_*τ*_)), is approximated by applying SLQ (Algorithm 2) to a special case of (5): We rewrite the log determinant as the trace of the matrix logarithm

#### Algorithm 3

Stochastic Lanczos derivativefree residual maximum likelihood (`SLDF-REML`)

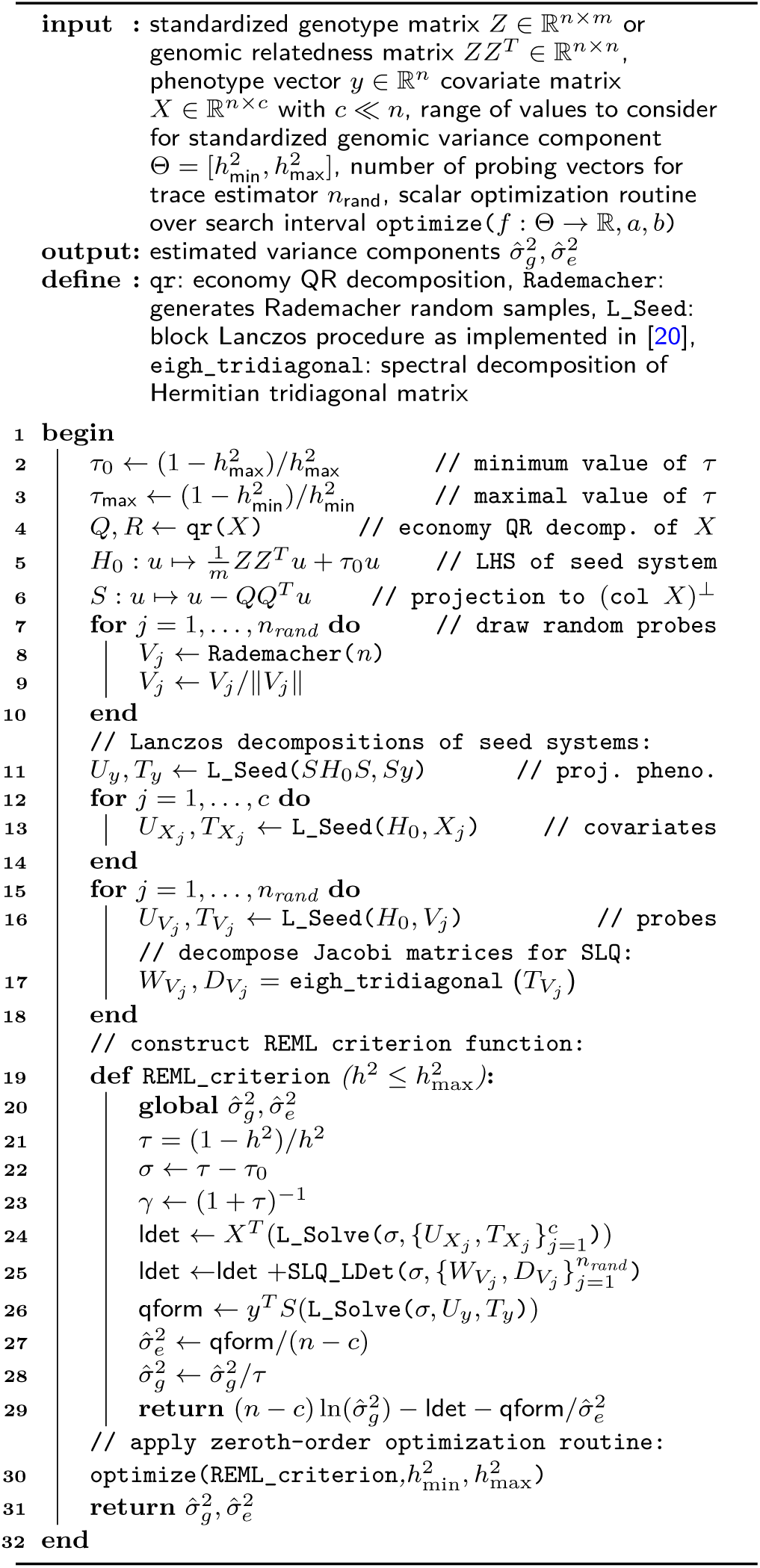

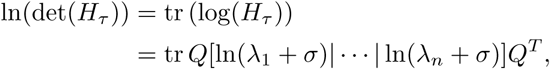

where we have spectrally decomposed *H*_0_ = *Q*Λ*Q*^*T*^ for some *τ*_0_ *≤ τ* with *σ* = *τ - τ*_0_. We draw *n*_rand_ *i.i.d.* normalized Rademacher random vectors 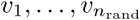 where each element of each vector *v*_*i*_ takes values of either 1/ ‖*v*_*i*_ ‖ or –1/ ‖*v*_*i*_ ‖ with equal probability. The SLQ approximate of the log determinant for the seed system is

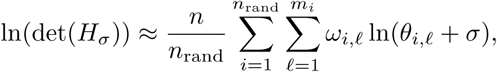

where the weights *w*_*i*,𝓁_ and nodes *θ*_*i*,𝓁_ are respectively derived by using the Lanczos process to construct orthonormal bases for *K*(*H*_0_, *v*_*i*_) (in practice, we apply block Lanczos to 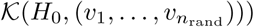) [16, 17].

### The SLDF_REML algorithm

Stochastic Lanczos derivative-free residual maximum likelihood (`SLDF_REML`; Algorithm 3), conceptually similar to the derivative-free algorithm of Graser and colleagues [8], applies the previously introduced Lanczos methods to approximate the above reparametrization of the REML criterion. Shift-invariance is then exploited such that, with the exception of the initial Lanczos decompositions, the REML log likelihood can be repeatedly evaluated using only vector operations. `SLDF_REML` takes a phenotype vector *y* ∈ ℝ^*n*^, a covariate matrix *X* ∈ ℝ^*n×c*^, either the genetic relatedness matrix *ZZ*^*T*^ ∈ ℝ ^*n×n*^ or the standardized genotype matrix *Z* ∈ ℝ^*n×m*^ (in which case the action of the GRM of a linear operator is coded implicitly as *v 1→ Z*(*Z*^*T*^ *v*)), and a range of possible standardized genomic variance component values 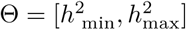 as arguments and generates a function `REML_criterion`: Θ *→ ℝ* that efficiently computes the log-likelihood of *τ |K*^*T*^ *y*. This function is then subject to scalar optimization via Brent’s method, which is feasible given the low cost of evaluation and low dimension of Θ. Hyperparameters include the number of probing vectors to be used for the SLQ approximation of the log determinant *n*_rand_, as well as tolerances corresponding to the REML criterion, parameter estimates, and the Lanczos residual norms. Convergence to a given tolerance on a sensible scale is ensured by optimizing over the standardized genomic variance component value *h*^2^ ∈ Θ ⊆ [0, 1] and evaluating the REML criterion at *τ* = (1 − *h*^2^)*/h*^2^. The REML criterion can be repeatedly evaluated in 𝒪 (*n*) operations, making high accuracy computationally feasible.

#### Algorithm 4

Lanczos first-order Monte Carlo residual maximum likelihood (`L_FOMC_REML`)

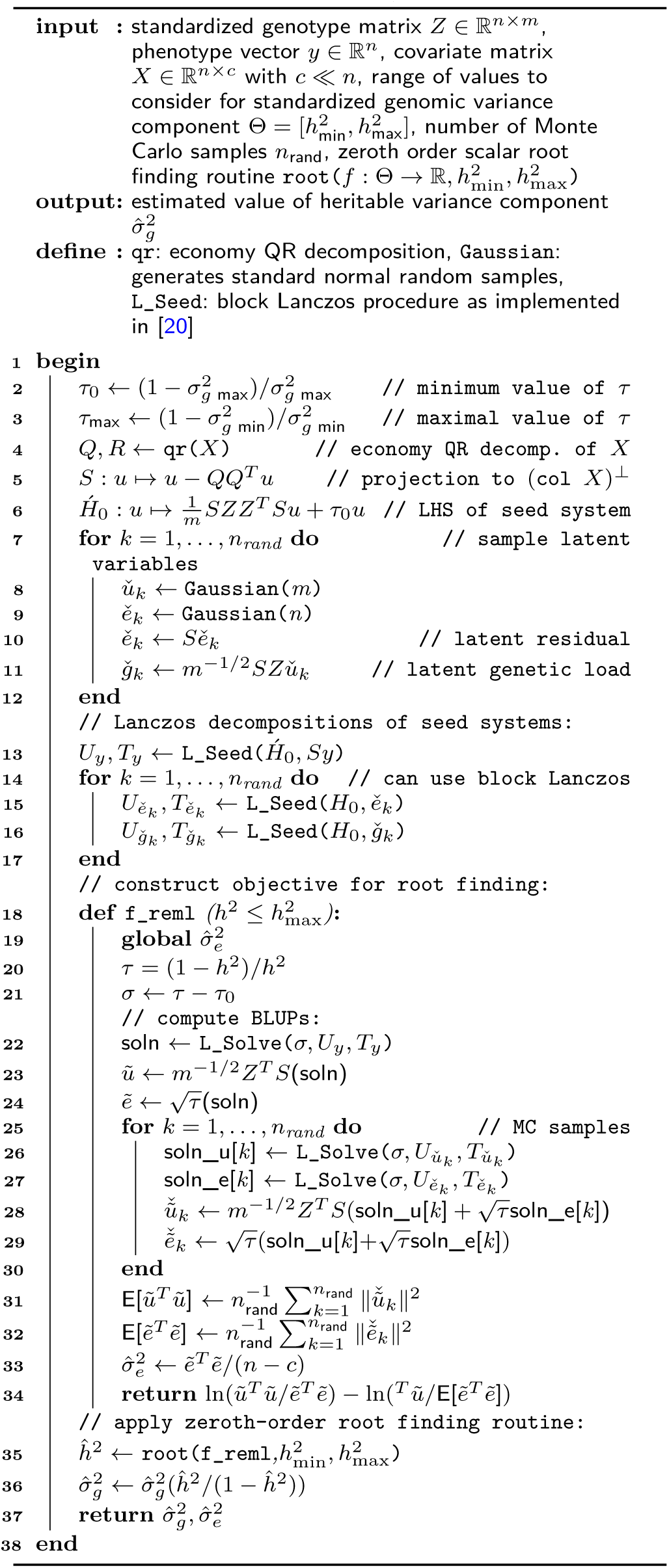

### A first-order Monte Carlo REML algorithm

We additionally propose the Lanczos first-order Monte Carlo residual maximum likelihood algorithm (`L_FOMC_REML`; Algorithm 4), which also takes advantage of the shifted structure of the standard genomic variance components model to speed computation. We first present the related first-order algorithm implemented in the efficient and widely-used BOLT-LMM software [5, 6], which we refer to as `BOLT_LMM` and of which the proposed `L_FOMC_REML` algorithm is a straightforward extension.

### BOLT_LMM (First-order Monte Carlo REML)

The `BOLT_LMM` algorithm is based on the observation that at stationary points of the REML criterion (3), the first order REML conditions (i.e., *∇𝓁* = 0) imply that

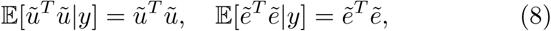

where ũ and 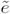 are the best linear unbiased predictions (BLUPs) of the latent genetic effects and residuals, respectively [22]. The BLUPs are functions of *τ* given by

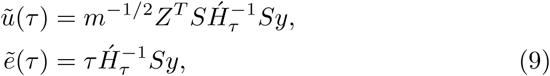

where we have defined 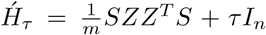 expectations (8) are approximated via the following stochastic procedure: Monte Carlo samples of the latent variables, 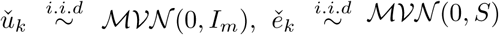 are used to generate samples of the projected phenotype vector

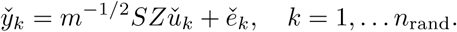

BLUPs are then computed as in (9), yielding the approximations

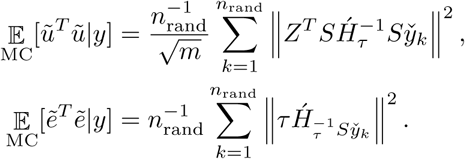

Using the above expressions, Loh et al. [5, 6] apply a zeroth-order root-finding algorithm to the quantity

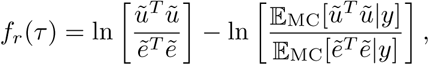

noting that *f*_*r*_ = 0 is a necessary condition (and, in practice, a sufficient condition) for (8). Using CG to approximate solutions to the linear systems involved in BLUP computations results in an efficient REML estimation procedure involving *O*(*n m n*_rand_) operations for well-conditioned covariance structures (i.e., for nontrivial non-heritable variance component values). As noted in [6], implicit preconditioning of *H*_0_ can be achieved by including the first few right singular vectors of the genotype matrix (or eigenvectors of the GRM) as columns of the covariate matrix *X*.

### The L_FOMC_REML algorithm

The `BOLT_LMM` algorithm described above involves solving *n*_rand_ + 1 linear systems

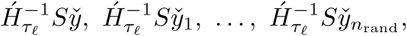

at each iteration of the optimization scheme in order to compute BLUPs of the latent variables for the observed phenotype vector and each of the Monte Carlo samples. However, each iteration involves spectral shifts of the left hand side of the form

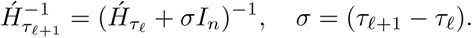

As in the `SLDF_REML` algorithm, the underlying block Krylov subspace is invariant to these shifts 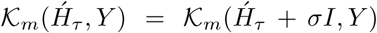. where 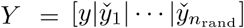 Thus, having performed the Lanczos process for an initial parameter value *τ*_0_, we can use `L_Solve` (Algorithm 1) to obtain the block CG ap-proximate solution 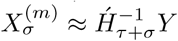 in *O* (*n* · *n*_rand_)erations. We are thus able to avoid solving linear systems in all subsequent iterations, though the relatively small number of matrix-vector products involved in computing BLUPs for the latent genetic effects at each step are unavoidable. The requirement of the genotype matrix for computing (9) prevents both `L_FOMC_REML` and `BOLT_LMM` from efficiently exploiting precomputed GRMs.

### Comparison of methods

We compare theoretical and empirical properties of our proposed algorithms, `SLDF_REML` and `L_FOMC_REML`, to those of `BOLT_LMM`.

### Computational complexity

In contrast to `BOLT_LMM`, the Lanczos-decomposition based algorithms we have proposed only need to perform the computationally demanding operations necessary to evaluate the REML criterion once. As such, we differentiate between *overhead* computations, which occur once and do not depend on the number of iterations needed to achieve convergence, and *per-iteration* computations, which are repeated until convergence of the optimization process (Table 1 and Fig. 4).

**Figure 4:**
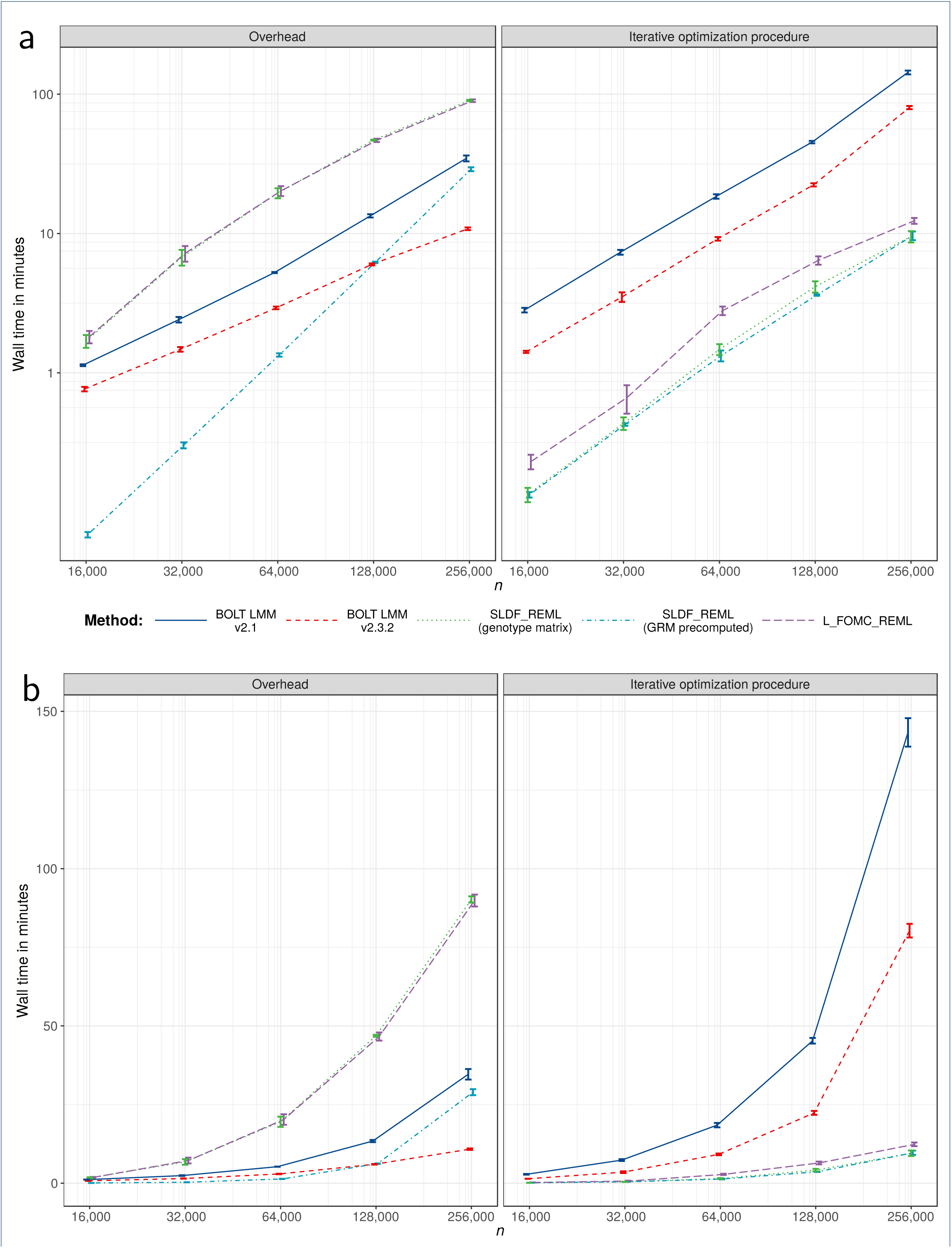
Overhead versus iterative optimization procedure timing results. Trimmed mean wall clock time for overhead computations and iterative REML optimization procedures across twenty replications per condition on the log_10_ scale (a) and natural scale (b). Error bars reflect per condition standard errors and lines connect per condition means.

The overhead computations of `SLDF_REML` are dominated by the need to construct bases for the *n*_rand_+*c*+1 subspaces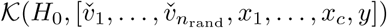, and are thus 𝒪 (*n*^2^(*n*_rand_ + *c*)*n*_*κ*_) when a precomputed GRM is available and 𝒪(2*m n*(*n*_rand_ + *c*)*n*_*κ*_) otherwise. Here, *n*_*κ*_ denotes the number of Lanczos iterations needed to achieve convergence at a pre-specified tolerance and increases with 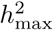. Subsequent iterations are dominated by the cost of solving *c* + 1 shifted linear systems via `L_Solve` and are thus 𝒪(*n · c · n*_*κ*_). The overhead computations in `L_FOMC_REML` are dominated by the Lanczos decompositions corresponding to the 2*n*_rand_ + 1 seed systems, where the GRM is implicitly represented in terms of the standardized genotype matrix, and is thus 𝒪 (4*m · n · n*_rand_ *n*_*κ*_). Operations of equivalent complexity are needed at *every* iteration of `BOLT_LMM`.

### Numerical experiments

We compared wall clock times for genomic variance component estimation for height in nested random subsets of 16,000, 32,000, 64,000, 128,000, and 256,000 unrelated 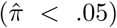 European ancestry individuals from the widely used UK Biobank data set [23]. All analyses included 24 covariates consisting of age, sex, and testing center and used hard-called genotypes from 330,723 array SNPs remaining after enforcing a 1% minor allele frequency cutoff. We compared `SLDF_REML`, with and without a precomputed GRM, to `L_FOMC_REML` which requires the genotype matrix. For the novel algorithms, absolute tolerances for the Lanczos iterations and the REML optimization procedure were set to 5e-5 and 1e-5, respectively. Additionally, we compared our interpreted Python 3.6 code to BOLT-LMM versions 2.1 and 2.3.3 (C++ code compiled against the Intel MKL and Boost libraries) [5, 6, 24, 25]. We ran each algorithm twenty times per condition, trimming away the two most extreme timings in each condition. Mirroring the default settings of the BOLT-LMM software packages, we set *n*_rand_ = 15 across both of our proposed methods.

Novel algorithms were implemented in the Python v3.6.5 computing environment [20], using NumPy v1.14.3 and SciPy v1.1.0 compiled against the Intel Math Kernel Library v2018.0.2 [25–27]. Optimization was performed using SciPy’s implementation of Brent’s method, with convergence determined via absolute tolerance of the standardized genomic variance component *ĥ* ^2^. Timing results (Table 2 and Figs. 3 and 5) do not include time required to read genotypes into memory, or, when applicable, to compute GRMs, and reflect total running time on an Intel(R) Xeon(R) Gold 6130 CPU @ 2.10GHz with 32 physical cores and 1 terabyte of RAM. Timing experiments excluded methods with cubic time complexity, including GCTA, FaST-LMM, and GEMMA. Accuracy was assessed by comparing heritability estimates generated by the stochastic algorithms to those estimated via the direct, deterministic average-information Newton– Raphson algorithm as implemented in the GCTA software package v1.92.0b2 [3] (Figures 2 and 5).

**Table 2:**
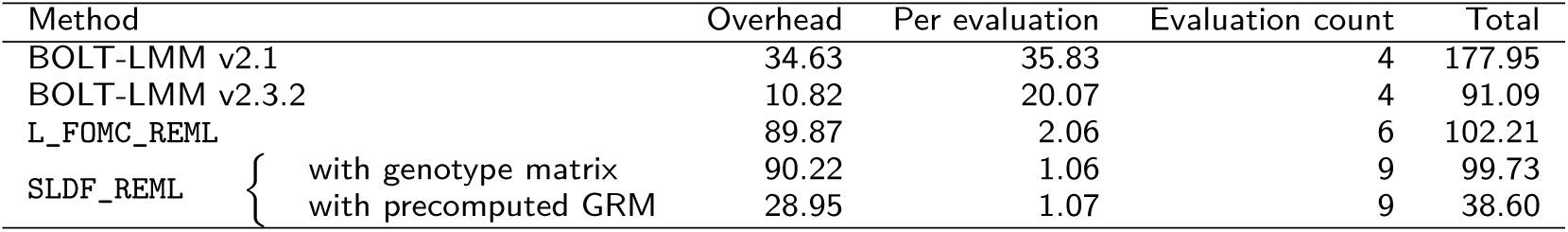
Overhead and per objective function evaluation timings of stochastic algorithms for *n*=256,000. Data reflect trimmed mean wall clock time in minutes over 20 iterations per condition.

**Figure 5:**
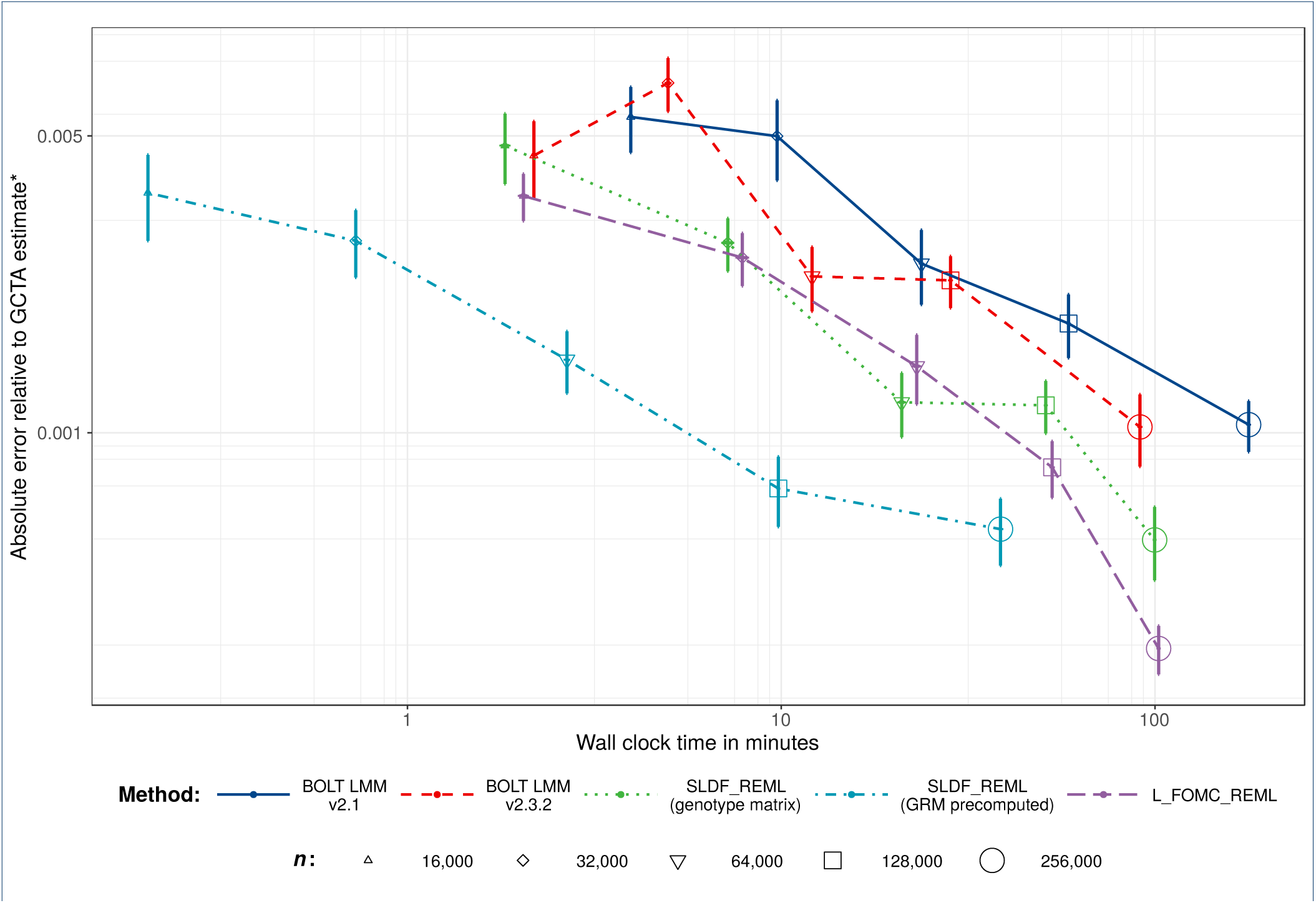
Numerical experiments: accuracy versus time. Average absolute error on the log_10_ scale with respect to the GCTA estimate^∗^versus trimmed mean wall clock time across twenty replications per condition. Standard Error bars reflect per condition standard errors and lines connect per condition trimmed means. ^*∗*^For *n*=256,000, memory requirements prohibited the use of GCTA, so we instead averaged ten estimates generated by the high-accuracy stochastic estimator implemented in BOLT-REML v2.3.2 [29] (standard errors were 6.32e-5 and 2.45e-7 for the mean heritability and its standard error, respectively).

## Results

Across 20 replications per condition for random subsamples of *n*=16,000 to 256,000 unrelated European-ancestry individuals, both `SLDF_REML` and `L_FOMC_REML` produced heritability estimates for height consistent with those generated by the GCTA software package (Figures 2 and 5). For large samples, the novel algorithms achieved greater accuracy than either version of BOLT-LMM. With respect to timings, `SLDF_REML` dramatically outperformed all other methods when the precomputed GRM was available (Table 2 and Fig. 3), which we expect whenever the number of markers exceeds the sample size. Examining methods taking genotype matrices as inputs, `SLDF_REML` and `L_FOMC_REML` performed similarly, whereas BOLT-LMM v2.3.2 converged more quickly than either in smaller samples (Figure 3), though the differences for *n*=256,000 were relatively minor (e.g., 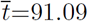 minutes for BOLT-LMM v2.3.2 versus 99.73 minutes for `SLDF_REML`; Table 2). The older version of BOLT-LMM, v2.1, performed significantly more slowly than any of the other implementations examined (e.g., average wall clock time was 177.95 minutes at *n*=256,000), demonstrating the importance of implementation optimization.

As the computations needed to compute the Lanczos decompositions in `L_FOMC_REML` and BOLT-LMM v2.3.2 are equivalent in time and memory complexity, we expect that an optimized compiled-language implementation of `L_FOMC_REML` would reduce the overhead computation time by a significant linear factor (*≈* 3 for *n*=256,000, comparing the sum of the overhead time and single objective function evaluation time for BOLT-LMM v2.3.2 to its total running time; Table 2). Consistent with theory, the wall clock times per objective function evaluation for the novel algorithms were trivial given the Lanczos decompositions (e.g., for *n*=256,000, 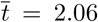 versus 20.07 minutes for `L_FOMC_REML` and BOLT-LMM v2.3.2, respectively; Table 2 and Fig. 4).

## Discussion

We have proposed stochastic algorithms for estimating the two component genomic variance component model (1), both of which theoretically offer substantial time savings relative to existing methods. Our methods capitalize on the principal of Krylov subspace shift invariance to reduce the number of steps involving 𝒪 (*n*^2^) or 𝒪 (*mn*) computations to one, whereas existing methods perform equivalent computations at each iteration of the REML optimization procedure. For large samples, when taking genotype matrices as inputs, our interpreted-language implementations of `L_FOMC_REML` and `SLDF_REML` [20] produced more accurate variance component estimates than the highly-optimized, compiled BOLT-LMM implementations, while taking similar amounts of time. Thus, we expect comparably-optimized implementations of the novel algorithms to compute high accuracy REML estimates in close to the time required by BOLT-LMM v2.3.2 for a *single* objective function evaluation. Further, in contrast to the `BOLT_LMM` algorithm, which requires the genotype matrix, `SLDF_REML` can exploit precomputed GRMs to reduce operation count by an 𝒪 (2*m/n*) factor (Table 1), which yields dramatic time savings when the number of markers greatly exceeds the number of individuals (Figure 3). While GRM precomputation is itself 𝒪 (*mn*^2^), it can be effectively and asynchronously parallellized across multiple compute nodes, substantially mitigating computational burden (though we note that serial input/output can interfere with efficient parallelization).

There are several limitations to the proposed approaches. First, `SLDF_REML`, which benefits from the ability to take GRMs as input, depends linearly on the number of included covariates, which might grow prohibitive in samples spanning numerous genotyping batches and ascertainment locations. However, as in `BOLT_LMM, L_FOMC_REML` requires 𝒪(*mn*) matrix multiplications for BLUP computation at each step of the REML optimization procedure, whereas `SLDF_REML` requires only vector operations. Thus, though the options provided by the two novel algorithms increase researchers’ flexibility overall, the choice of whether to employ `SLDF_REML` versus `L_FOMC_REML` is problemspecific and necessitates greater researcher attention to resource allocation. Second, neither algorithm mitigates the substantial *𝒪* (*mn*) or *𝒪* (*n*^2^) memory complexity common to all algorithms for REML estimation of genomic variance components, requiring that researchers have access to high-memory compute nodes to work with large samples (though we note that neither of the novel algorithms substantial increases this burden either). Finally, for the same reasons that the spectral decomposition-based direct methods implemented in the FaST-LMM and GEMMA packages [2, 4] are restricted to the simple two component model (i.e., whereas the GRM and identity matrix are simultaneously diagonalizable, the same doesn’t hold for arbitrary collections of three or more symmetric positive semidefinite matrices), the shift-invariance property exploited by the proposed methods does not extend to multiple genomic variance components. Given that the two component model is insufficient for precise heritability estimation for many complex traits [28], our novel algorithms apply to the singular, though common, task of variance component estimation for LMM in association studies.

Despite these limitations, the proposed algorithms have clear advantages over existing methods in terms of flexibility, accuracy, and speed of computation. We provide both pseudocode and heavily annotated Python 3 implementations [20] to facilitate their incorporation into existing software packages. Finally, we suggest that the methods presented in our theoretical development, in particular stochastic trace estimation and stochastic Lanczos quadrature, are likely to find uses in REML estimation of other models of interest to researchers in genomics.

## Abbreviations

BLAS: Basic Linear Algebra Subprogram
BLUP: Basic Linear Unbiased Prediction
CG: Conjugate Gradients method
GCTA: Genome-wide Complex Trait Analysis [3]
GRM: Genomic Relatedness Matrix
GWAS: Genome-Wide Association Study
LMM: Linear Mixed-effects Model
REML: Residual Maximum Likelihood
SLQ: Stochastic Lanczos Quadrature

## Declarations

Ethics approval and consent to participate

UK Biobank data collection procedures were approved by the UK Biobank Research Ethics Committee (reference 11/NW/0382).

### Consent for publication

Not applicable.

### Availability of data and material

The UK Biobank data are available to qualified researchers via the UK Biobank Access Management System (https://bbams.ndph.ox.ac.uk/ams). The code used in the numerical experiments is available on Github (https://github.com/rborder/SL_REML).

### Authors’ contributions

RB wrote the manuscript, developed the algorithms, wrote the code used in numerical experiments, and analyzed the data. SB supervised the project and contributed to the development of the algorithms and the writing of the manuscript.

## Acknowledgements

The authors wish to thank UK Biobank participants. Additionally, the authors thank Matthew C. Keller and Luke M. Evans for their thoughtful comments and provision of computational resources.

## Additional Files

Additional file 1 — Appendix.pdf

Proof of result used to efficiently compute the quadratic form (7).

