## Appendix for "Stochastic Lanczos estimation of genomic variance components for linear mixed-effects models"

Let $A^+$ denote any generalized inverse and let $A^\dagger$ denote the Moore-Penrose pseudoinverse.

**Fact 1** (Seber [2008], page 457)**.** *Let $S \in \mathbb{R}^{n\times n}$ be an orthogonal projection operator. Then for any $B \in \mathbb{R}^{m\times n}$,*

$$S(BS)^\dagger = (BS)^\dagger$$

*and*

$$(SB)^\dagger S = (SB)^\dagger$$

**Fact 2** (Lewis and Newman [1968], Theorem 6)**.** *Let $A \in \mathbb{R}^{n\times n}$ be such that $A = A^T \succeq 0$, $C \in \mathbb{R}^{n\times c}$, and define $\tilde{A} = A + C^T C$. Let $A^+ \succeq 0$ be a generalized inverse of $A$. Then*

$$\tilde{A}^+ = A^+ - A^+ C^T (I + CA^+ C^T)^{-1} CA^+$$

*is a generalized inverse of $\tilde{A}$ if and only if $\ker A \subseteq \ker C$. Further, if this is the case, we have*

$$\begin{aligned}
\tilde{A}^+ \tilde{A} &= A^+ A, \\
\tilde{A}\tilde{A}^+ &= AA^+, \\
\tilde{A}^+ &\succeq 0, \\
\text{and } \tilde{A}^+ = \tilde{A}^\dagger &\iff A^+ = A^\dagger.
\end{aligned}$$

**Lemma 1.** *Write the full QR decomposition of $X \in \mathbb{R}^{n\times c}$ as $X = QR = [Q_X | Q_{X^\perp}]R$ such that $Q_X \in \mathbb{R}^{n\times(n-c)}$, $Q_{X^\perp} \in \mathbb{R}^{n\times c}$ and define $S = I_n - Q_X Q_X^T$. Then,*

$$y^T S(S(H+\sigma I)S)^\dagger Sy = y^T S(SHS + \sigma I)^{-1} Sy$$

*Proof.* Applying the first fact,

$$y^T S(S(H+\sigma I)S)^\dagger Sy = y^T (S(H+\sigma I)S)^\dagger y.$$

Rewrite the left hand side as

$$\begin{aligned}
y^T (S(H+\sigma I)S)^\dagger y &= \sigma^{-1} y^T (\underbrace{\sigma^{-1} SHS}_{\equiv A} + SS)^\dagger y \\
&= \sigma^{-1} y^T \tilde{A}^\dagger y,
\end{aligned}$$

where we define $\tilde{A} = A + S^T S$. Note that the first fact also yields

$$SA^\dagger S = SA^\dagger = A^\dagger S = A^\dagger$$

Using the second fact, we then have

$$\begin{aligned}
\sigma^{-1} y^T \tilde{A}^\dagger y &= \sigma^{-1} y^T (A^\dagger - A^\dagger S(I + SA^\dagger S)^{-1} SA^\dagger) y \\
&= \sigma^{-1} y^T (A^\dagger - A^\dagger (I + A^\dagger)^{-1} A^\dagger) y \\
&= \sigma^{-1} y^T (A + I)^{-1} y \\
&= y^T S(SHS + \sigma I)^{-1} Sy
\end{aligned}$$

□
